# Fertility-LightGBM: A fertility-related protein prediction model by multi-information fusion and light gradient boosting machine

**DOI:** 10.1101/2020.08.24.264325

**Authors:** Lingling Yue, Minghui Wang, Xinhua Yang, Yu Han, Lili Song, Bin Yu

**Affiliations:** College of Mathematics and Physics, Qingdao University of Science and Technology, Qingdao 266061, China; Artificial Intelligence and Biomedical Big Data Research Center, Qingdao University of Science and Technology, Qingdao 266061, China; School of Life Sciences, University of Science and Technology of China, Hefei 230027, China

**Keywords:** Fertility-related protein, Multi-information fusion, LASSO, LightGBM

## Abstract

The identification of fertility-related proteins plays an essential part in understanding the embryogenesis of germ cell development. Since the traditional experimental methods are expensive and time-consuming to identify fertility-related proteins, the purposes of predicting protein functions from amino acid sequences appeared. In this paper, we propose a fertility-related protein prediction model. Firstly, the model combines protein physicochemical property information, evolutionary information and sequence information to construct the initial feature space ‘ALL’. Then, the least absolute shrinkage and selection operator (LASSO) is used to remove redundant features. Finally, light gradient boosting machine (LightGBM) is used as a classifier to predict. The 5-fold cross-validation accuracy of the training dataset is 88.5%, and the independent accuracy of the training dataset is 91.5%. The results show that our model is more competitive for the prediction of fertility-related proteins, which is helpful for the study of fertility diseases and related drug targets.

## 1. Introduction

In the early stages of development, fertility-related proteins participate in many aspects of life activities [1]. It not only plays a regulatory role in complex fertility-related events [2, 3] but also plays a crucial role in many biological entities [4, 5]. The identification of fertility-related proteins is helpful to decipher the potential mechanism of fertility-related events, and then to understand their molecular functions in detail, to provide a theoretical basis for the development of related drugs.

Park et al. [6] used two-dimensional electrophoresis and western blot analysis to study the relationship between protein expression and bull physical characteristics. To understand the source of sperm heterogeneity, D’ Amours et al. [7] extracted protein from low-density and high-density sperm by Percoll gradient centrifugation and sodium deoxycholate, and analysed proteomics by isobaric tag for relative and absolute quantitation. Schumacher et al. [8] studied the possible connection between mammalian sperm protein sequence evolution and human phosphorylation status, where immunoblotting, mass spectrometry and two-dimensional gel electrophoresis were combined to identify 99 sperm proteins. Moura et al. [9] evaluated the protein expression in accessory sex gland fluid and its relationship with the reproductive index of dairy cows. Chen et al. [10] provided system-level insights into sexual dimorphism and gametogenesis through gene ontology annotation and path analysis. Kwon et al. [11] used proteomics to reduce the energy of boar sperm, and they constructed related signaling pathways based on differentially expressed proteins to identify proteins associated with sperm capacitation. Légaré et al. [12] used differential proteomics with isobaric tags for relative and absolute quantitative labeling. They used liquid chromatograph-mass spectrometer analysis to identify fertile and infertile male sperm differentially expressed proteins.

Limited by the complex protein functions and experimental process, it may take months or longer to determine the protein functions. For this reason, researchers continually develop computational models to predict protein functions from amino acid sequences. Rahimi et al. [13] developed OOgenesis to identify oogenesis-related proteins by six feature extraction methods and the support vector machine (SVM). However, this model, which has certain limitations, can only identify a protein related to fertility. Therefore, Bakhtiarizadeh et al. [14] constructed the first general model PrESOgenesis for predicting fertility proteins, which trained a two-layer classification model based on the SVM, and the first layer identified whether the protein is related to fertility. The second layer determined what kind of fertility is associated with this protein. PrESOgenesis can achieve 82.97% accuracy and still has promotion space. On this basis, Le [15] proposed the Fertility-GRU method to distinguish fertility-related proteins. By employing the gated recurrent unit (GRU) architecture, Fertility-GRU saved the position-specific scoring matrix (PSSM) information into a deep neural network to prevent the loss of sequence information as much as possible, and achieved 91.1% prediction accuracy on the independent test dataset. However, data and features determine the upper limit of machine learning. The feature vector obtained by the single feature extraction method is too monotonous to express the information of the protein related to fertility fully. Based on this, we propose a new prediction model to identify fertility-related proteins.

Our model is a prediction model which is suitable for general fertility-related proteins and considers the sequence information, physicochemical property information and evolutionary information. Firstly, we choose pseudo position-specific scoring matrix (PsePSSM), amino acid composition (AAC), dipeptide composition (DC), composition transition distribution (CTD), autocorrelation descriptor (AD) and encoding based on grouped weigh (EBGW) to extract amino acid residue information, then we fuse the feature vectors. Secondly, we use LASSO to eliminate the redundant features and retain useful features. Finally, LightGBM is used for classification, and the prediction results are compared with the existing models.

## 2. Materials and methods

### 2.1. Datasets

The effectiveness of statistical forecasting tools depends on the availability of high-quality data. Training data need to be accurate, organized and as complete as possible to maximize predictability. Bakhtiarizadeh et al. [14] created a protein initial positive dataset by searching the UniProt Knowledgebase (UniProtKB) and rejected proteins with sequences higher than 6000 or less than 60. Then they deleted paired sequences with similarity higher than 50% in the same subset by CD-HIT program [16] and removed protein sequences that contain ambiguous residues (‘B’, ‘X’ or ‘Z’). On this basis, the redundant sequences were deleted, and 1704 fertility-related proteins were finally obtained. In the same way, Le [15] provided 1815 non-fertility-related proteins.

In this paper, we randomly divide the above two kinds of proteins into the training dataset *Strain* and the independent test dataset *S*_*test*_. The relevant sets are defined as follows:

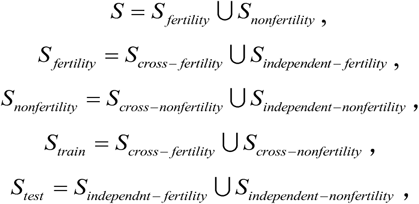

where *S* represents the protein dataset used in this paper, which is composed of *S* _*fertility*_ (including 1704 fertility-related proteins) and *S*_*nonfertility*_ (including 1815 non-fertility-related proteins). *S*_*cross − fertility*_ is a set consisting of 1420 fertility-related proteins randomly taken from *S* _*fertility*_. The remaining 284 fertility-related proteins in *S* _*fertility*_ are recorded as *S*_*indepengdent* − *fertility*_. *S*_*cross* − *nonfertility*_ is a set consisting of 1512 non-fertility-related proteins randomly taken from *S*_*nonfertility*_. The remaining 303 non-fertility-related proteins in *S*_*nonfertility*_ are marked as *S*_*independent* − *nonfertility*_.

### 2.2. Feature extraction

Feature coding, which can convert protein sequence information into numerical information, is a critical step in building a classification model. We use the following six feature coding methods.

#### 2.2.1. Pseudo position-specific scoring matrix

Pseudo position-specific scoring matrix (PsePSSM) proposed by Chou and Shen [17] is widely used in proteomics prediction [18-21]. We use the PSI-BLAST program [22] to perform three iterative searches with *E* value of 0.001 for UniProtKB / Swiss-Prot database. The PSSM [23] matrix corresponding to each protein sequence is obtained as follows:

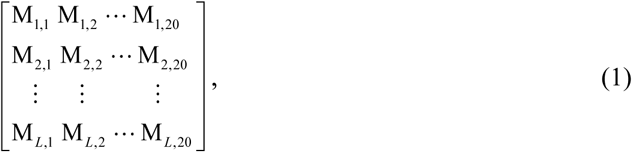

where *L* is the length of *P*, M_*i, j*_ (*i* = 1, 2, …, *L*; *j* = 1, 2, …, 20) is the position-specific score obtained by mutation of amino acid residue at location *i* to residue *j* during evolution. In order to reduce the deviation,

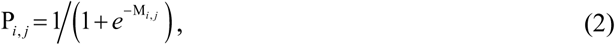

we normalize M_*i, j*_ to P_*i, j*_ based on (2), P_*i, j*_ ∈ (0,1) and then convert (1) into

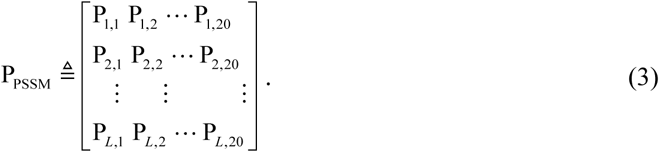

Since the lengths of protein sequences in the dataset are not same, it is necessary to transform protein sequences into a vector with uniform dimensions using the following formula:

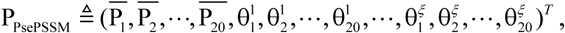

where

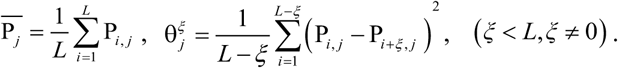

#### 2.2.2. Amino acid composition

The amino acid composition (AAC) widely used in proteomic research [24, 25] was proposed by Nakashima and Nishikawa [26]. This method calculates the frequency of 20 amino acids on each protein. Each protein sequence *P* can be represented by vector (*v*_1_, *v*_2_, …, *v*_20_)^*T*^ through AAC, that is,

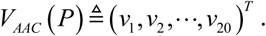

*v*_*i*_, which can be calculated by *v*_*i*_ = *f*_*i*_ */L*, is the frequency of the *i* amino acid.

#### 2.2.3. Dipeptide composition

Dipeptide composition (DC) [27-29] calculates the frequency of dipeptide (amino acid pair). DC not only considers the coupling between two neighboring residues, but also can adequately reflect the composition and sequence information of amino acids. Twenty amino acids constitute 20 × 20 = 400 amino acid pairs. Therefore,

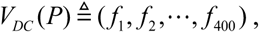

where *f*_*i*_ is the frequency of the *i* amino acid pair in sequence *P*.

#### 2.2.4. Composition transition distribution

Composition transition distribution (CTD) [30, 31] can replace amino acid residues with their class indexes. First of all, as shown in Fig. S1, we divide each protein sequence into ten segments with different lengths and groups in order to describe the continuous and discontinuous interaction patterns of multiple overlapping residues. Furthermore, we divide amino acids according to dipole and side-chain volume for reducing the internal complexity of amino acids and adapting to synonymous mutation of amino acids. The grouping is shown in Table S1. For each local fragment region, we calculate the following three descriptors.

(1) Composition (C) is calculated by

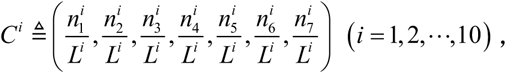

where *C*^*i*^ and *L*^*i*^ represent component descriptor and length of sequence for local fragment region *i*, respectively. 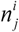 is the number of times that seven groups *j* of amino acids appear in the local fragment *i*.

(2) Transition (T) is the frequency of dipeptides that can be composed of seven groups of amino acids, which is calculated by

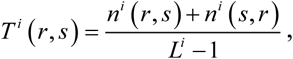

where *n*^*i*^ (*r, s*) and *n*^*i*^ (*s, r*) represent the number of occurrences of the amino acid pair (*r, s*) and (*s, r*) in the *i*, respectively. Each local segment produces 21 features.

(3) Distribution (D) represents the distribution pattern of each group of amino acids. This distribution pattern is measured sequentially along the first, 25%, 50%, 75%, and 100% positions of each group. It can be calculated as

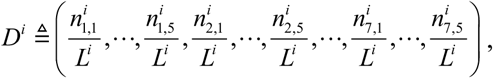

where 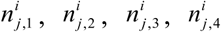, and 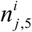 are five descriptors of distribution for every attribute in first residue, 25% residue, 50% residue, 75%residue, and 100% residue, respectively.

#### 2.2.5. Autocorrelation descriptors

Autocorrelation descriptors (AD) are defined according to the distribution of amino acids along the sequences, which are widely used in proteomics research [32]. 566 amino acid indices were collected in the amino acid index (AAindex) database [33] of version 9.2. We select seven amino acid indices as shown in Table S2. Due to the different measurement units of various physicochemical properties, all indicators need to be centralized and standardized by

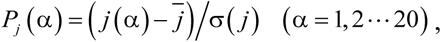

where *P*_*j*_ (*α*) is the *j* physicochemical property index of the *α* amino acid after linear transformation, *j* (*α*) is the original index of *α*. 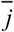 and *σ* (*j*) represent the mean and standard deviation of the physicochemical properties for *j*, respectively. On this basis, the following three descriptors are used to convert protein letter sequences into digital signals.

The Moran autocorrelation descriptor is thus defined as:

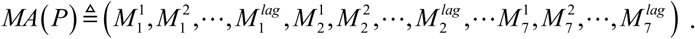

*L* represents the length of *P, lag* represents a built-in parameter, which represents the autocorrelation lag interval. Each element 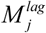 can be calculated by (4):

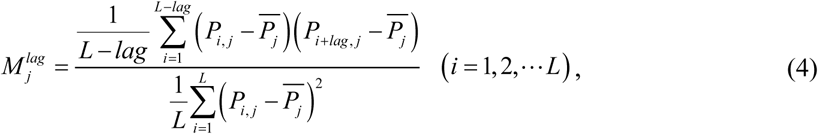

where *P*_*i, j*_ is the corresponding value of the *j* index at the *i* -th position in *P*, 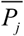 represents the mean of the *j* index in *P*.

The Geary autocorrelation descriptor is defined as follows:

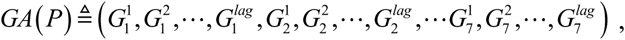

where *GA*(*P*) is the Geary autocorrelation factor of *P*. Each element 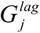 can be calculated by (5):

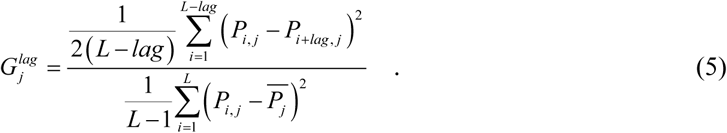

The normalized Moreau-Broto autocorrelation descriptor is defined as:

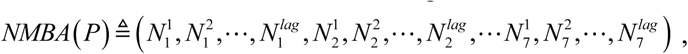

where *NMBA*(*P*) is the normalized Moreau-Broto autocorrelation factor of *P*. Each element 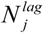 can be calculated by (6):

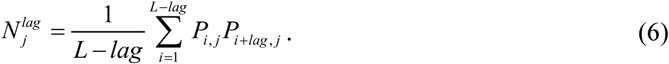

#### 2.2.6. Encoding based on grouped weigh

Zhang et al. [34] proposed an encoding based on grouped weigh (EBGW) that can effectively extract the physicochemical property information of proteins [35-38].

Amino acids are divided into four groups according to their physicochemical properties, and three new partition methods are obtained by combining two non-intersect groups. The detailed introduction is shown in Supplementary Si1. Each protein sequence *P* = *α*_1_*α*_2_ … *α*_*n*_ is mapped to three binary sequences of length *n* :

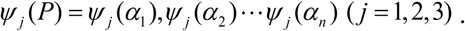

*ψ* _*j*_ (*α*_*i*_) (*i* = 1, 2, …, *n*) is calculated by (7), (8) and (9).

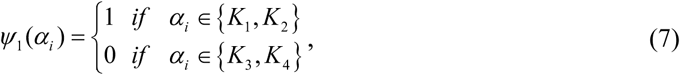

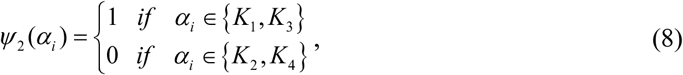

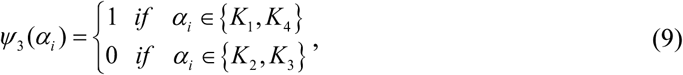

where *α*_*i*_ (*i* = 1, 2, …, *n*) represents the *n* -th amino acid in *P*. Then each binary sequence is divided into *L* subsequences. The normal weight of the *l* -th subsequence of the *j* -th binary sequence is

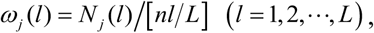

where *N* _*j*_ (*l*) and [*nl L*] are the number of occurrences of ‘1’ and the length of the subsequence, respectively. [·] is the rounding operation. Thus, each protein sequence *P* can be recorded as:

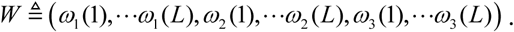

### 2.3. Feature selection

In order to reduce the fitting risk, the compression estimation method LASSO [19, 39, 40] can add penalty terms to the coefficients on the basis of least squares. The features with the small contribution of the model are removed.

Supposed the dataset *D* = {(*x*_1_, *y*_1_),(*x*_2_, *y*_2_),… …, (*x*_*m*_, *y*_*m*_)}, where *x* ∈ *R*^*d*^, *y* ∈ *R*, the optimization goal is as follows:

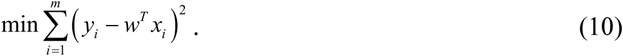

Eq. (10) is ordinary linear regression. In order to reduce the risk of overfitting, we use LASSO and introduce *𝓁*_1_ -norm regularization on the basis of the minimum residual square sum:

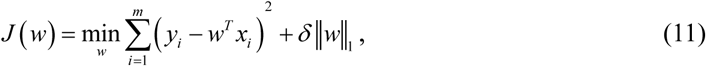

The detailed introduction is shown in Supplementary Si2.

### 2.4. Machine learning

LightGBM [41-43] is an improvement of the gradient boosting decision tree (GBDT) algorithm [44], and its core principle is based on the decision tree algorithm. It builds decision trees by leaf-growing strategies. Limiting the maximum depth of trees can not only ensure the training efficiency but also prevent overfitting. The algorithm introduces the techniques of gradient-based one-side sampling (GOSS) and exclusive feature bundling (EFB) into the traditional GBDT algorithm.

The detailed introduction of GOSS is shown in Supplementary Si3. Firstly, the absolute values of the gradients of the training examples are sorted in descending order, and the data with the gradient value before *a* × 100% is selected as the set *A*. Secondly, in the remaining instance *A*^*c*^, a subset *b* × |*A*^*c*^| of size *B* is randomly chose. Finally, the variance gain of the set *A* ∪ *B* is calculated according to the formula

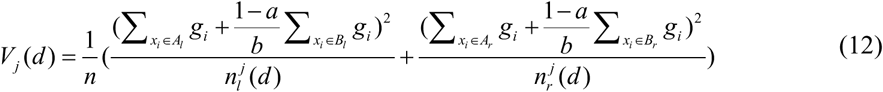

to segment the instance, where *A*_*l*_ = {*x*_*i*_ ∈ *A*:*x*_*ij*_ ≤ *d*}, *A*_*r*_ = {*x*_*i*_ ∈ *A*:*x*_*ij*_ > *d*}, *B*_*l*_ = {*x*_*i*_ ∈ *B*:*x*_*ij*_ ≥ *d*}, *B*_*r*_ = {*x*_*i*_ ∈ *B*:*x*_*ij*_ < *d*}. *g*_*i*_ and *d* are the gradient of sample *i* and the segmentation point of segmentation feature, respectively. 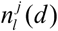 and 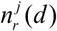 represent the number of samples whose value is less and greater than or equal to *d* on the *j* -th feature, respectively.

The high-dimensional features are usually sparse, and many features are mutually exclusive in the sparse features. In order to reduce the number of features, EFB is used to bundle exclusive features. EFB binds mutually exclusive features as a single feature by carefully designing feature scanning algorithm. The complexity of constructing the histogram is reduced from O(*data* * *feature*) to O(*data* * *bundle*) in this way. Due to *bundle ≪ feature*, the amount of features to be traversed is greatly reduced. By this method, the training process is greatly accelerated without the loss of features.

### 2.5. Model evaluation

In this paper, 5-fold cross-validation and independent dataset test are used to derive comparative metrics (values) amongst the reviewed predictors. Sensitivity (Sen), specificity (Spe), accuracy (Acc) and Matthew’s correlation coefficient (MCC) are used as evaluation indicators. The above indicators are defined as follows:

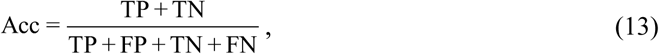

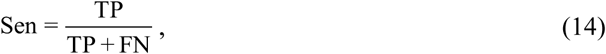

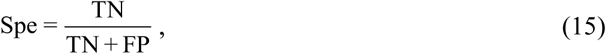

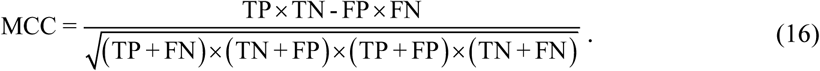

In addition, the area under curve (AUC) and the area under precision recall (AUPR) are also important indicators for measuring the robustness of the model. The AUC value and AUPR are the sizes of the area under the receiver operating characteristic (ROC) curve and the precision recall (PR) curve, respectively. The detailed introduction is shown in Supplementary Si4.

### 2.6. Our model: Fertility-LightGBM

Fertility-related proteins prediction models have been proposed in many papers. PrESOgenesis [14] was a two-tier classification model based on SVM. The first tier can classify fertility-related proteins and non-fertility-related proteins, and the second tier can identify proteins related to oogenesis, spermatogenesis and embryogenesis. In PrESOgenesis, SVM was used as a classifier, and radial basis function was selected. Radial basis function is too dependent on parameters, and it often takes too long to meet performance requirements when facing large-scale training samples. LightGBM supports parallel learning, which can process massive data and has higher learning efficiency. Fertility-GRU [15] saved all PSSM information to the convolutional neural network for prediction through GRU architecture. But a single feature extraction method often gets too monotonous information to represent fertility-related protein features. The method of multi-information fusion can fully consider protein sequence features. In conclusion, we propose Fertility-LightGBM.

Fig. 1 shows the specific steps of Fertility-LightGBM:

***Step 1:*** The dataset of fertility-related proteins is obtained, and input the protein sequences and their corresponding binary classification problem class labels.

***Step 2:*** Feature extraction. Transform protein sequence signals into numerical signals through PsePSSM, AAC, DC, CTD, AD and EBGW methods. The initial feature space is constructed by fusing the feature vectors end to end.

***Step 3:*** Feature selection. LASSO is used to remove the redundant information while retaining the essential classification features to choose the optimal feature subset.

***Step 4:***The best feature subset and the true labels are input into the LightGBM for prediction according to ***Step 2*** and ***Step 3***.

***Step 5:*** Model evaluation. Sen, Spe, Acc, MCC, AUC and AUPR are used to evaluate the predictive performance of Fertility-LightGBM. Then test the generalization ability by independent dataset test.

**Fig. 1.**
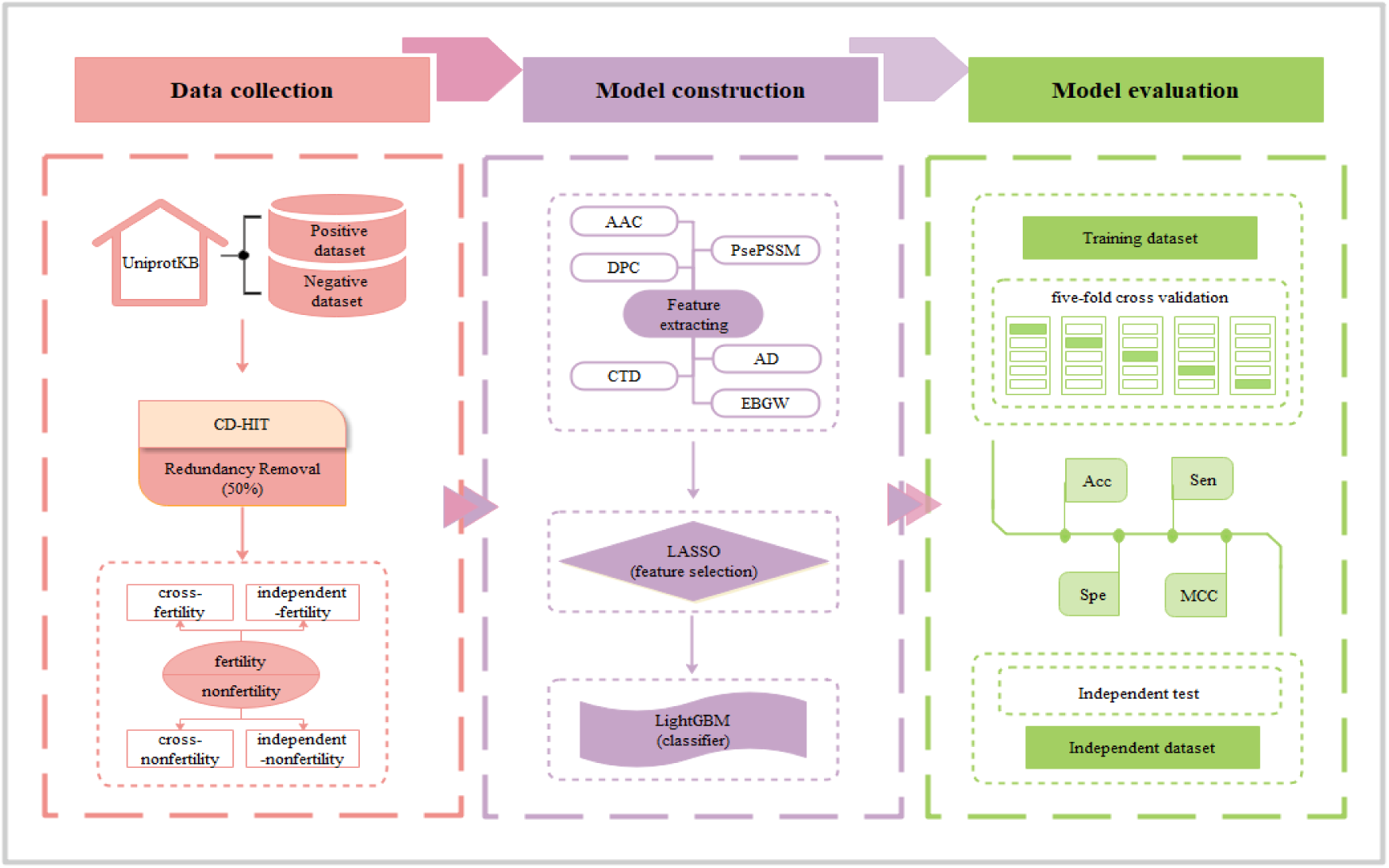
The framework of Fertility-LightGBM.

## 3. Results and discussion

### 3.1. Parameter selection of feature extraction algorithm

It is necessary to determine the best parameters *ξ, lag* and *L* of PsePSSM, AD and EBGW in feature extraction for the prediction ability of our model. Limited by the sequence length, the parameters of PsePSSM are set from 1 to 50. For determining the optimal parameters, we choose LightGBM and 5-fold cross-validation. Acc (the most important metrics), Sen, Spe and MCC are evaluation indicators. The prediction results corresponding to different parameters on the training dataset are shown in Fig. 2. Table S3-S5 shows specific prediction results.

**Fig. 2.**
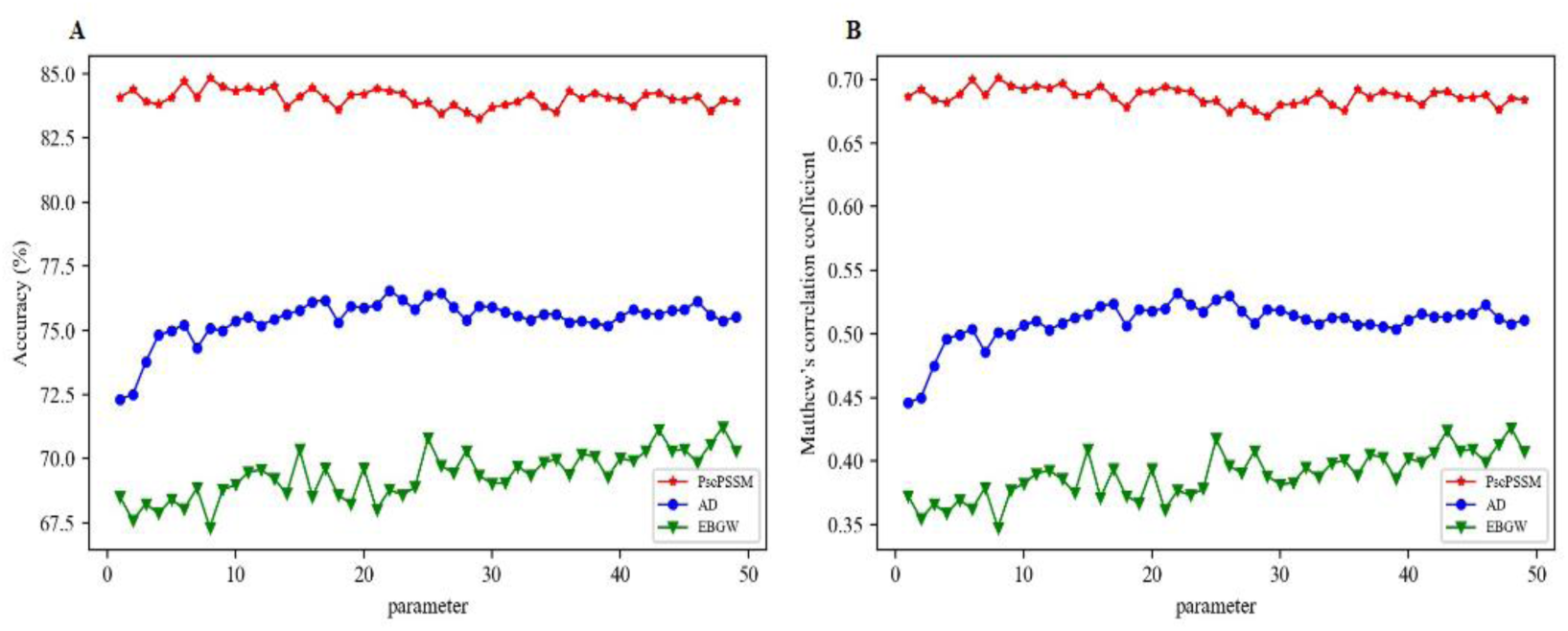
The prediction results of various parameters on the training dataset.(A) Acc predicted by different parameters of PsePSSM, AD and EBGW. (B) MCC predicted by different parameters of PsePSSM, AD and EBGW.

Fig. 2 shows that the different values of *ξ* in PsePSSM algorithm will change the prediction results on the training dataset. When *ξ* =9, Acc and MCC reach 84.83% and 0.7009 respectively, which are 0.10%-1.57% and 0.10%-2.99% higher than those predicted by other parameters. We analyze the model prediction performance when *ξ* takes different values. The best parameter of PsePSSM is 9.

We use three descriptors and seven amino acid indexes so that each protein sequence can be represented by an 3 × 7 × *lag* -dimensional vector, where *lag* represents the built-in parameter of AD encoding. Fig. 2 shows that the prediction results are very different for the different *lag*. When *lag* takes 23, Acc can reach a maximum of 76.55%, and the MCC also reaches a maximum value of 0.5320. Therefore, the optimal parameter of the AD algorithm is 23. Each protein generates a 3× 7 × 23 = 483 -dimensional feature vector by AD.

Fig. 2 shows that prediction results vary with the number of sub-sequences *L*. When the value of *L* is 49, both Acc and MCC can reach maximum values of 71.23% and 0.4260, respectively, which are 0.10% - 4.6% and 0.23% - 9.16% higher than others. The prediction effect is the best when the number of subsequences is 49.

### 3.2. Influence of feature extraction methods

Feature extraction methods can digitize the protein letter sequences and express them in the form of feature vectors, which can reflect the intrinsic correlation between the sequence and the expected target. We extract features from proteins by six feature codes. The extracted feature information is connected end-to-end according to the sequence of PsePSSM, AAC, DC, CTD, AD and EBGW, then the 1880-dimensional initial feature space ‘ALL’ is obtained. We use LightGBM as a classifier, and get the prediction results through 5-fold cross-validation evaluation model on the training dataset. The results of different feature extraction methods are shown in Table 1.

**Table 1.** The prediction results of different features on the training dataset.

| Feature space | Acc (%) | Sen (%) | Spe (%) | MCC |
| --- | --- | --- | --- | --- |
| PsePSSM | 84.83 | 87.04 | 82.76 | 0.7009 |
| AAC | 79.86 | 80.14 | 79.59 | 0.5982 |
| DC | 79.04 | 75.28 | 82.56 | 0.5820 |
| CTD | 76.21 | 75.21 | 77.14 | 0.6255 |
| AD | 76.55 | 72.04 | 80.78 | 0.5320 |
| EBGW | 71.23 | 72.25 | 70.28 | 0.4260 |
| ALL | **88.07** | 88.24 | 87.91 | **0.7640** |

Table 1 shows that the Acc and MCC of the ‘ALL’ are 88.07% and 0.7640, which are 3.24%-16.84% and 6.31%-33.80% higher than those of any single feature extraction method. It indicates that a single feature extraction method limits the prediction ability, and the fusion feature method can obtain more effective biological feature information from the protein sequences. Therefore, we adopt the technique of multi-information fusion.

### 3.3. Influence of feature selection methods

Multi-information fusion will produce more redundant features while increasing the storage requirements and computational costs of data analysis. Reducing the dimensions is necessary to construct an ideal fertility-related proteins prediction model. In order to mine important features from high-dimensional data and improve model robustness, we adopt mutual information (MI) [45], factor analysis (FA) [46], kernel principal component analysis (KPCA) [47], locally linear embedding (LLE) [48], principal component analysis (PCA) [49], truncated singular value decomposition (TSVD) [50], spectral embedding (SE) [51] and LASSO to reduce the dimensions. For comparing eight feature selection methods and choosing the best feature subset, the different subsets of features that filter by different methods are used to input the LightGBM, then test these results by 5-fold cross-validation. The prediction results of different feature selection methods on the training dataset are shown in Table 2 and Fig. 3.

**Table 2.** The prediction results of eight feature selection methods on the training dataset.

| Method | Dimensions | Acc (%) | Sen (%) | Spe (%) | MCC |
| --- | --- | --- | --- | --- | --- |
| MI | 57 | 84.63 | 86.20 | 83.15 | 0.6980 |
| FA | 57 | 85.28 | 84.01 | 86.46 | 0.7072 |
| KPCA | 57 | 85.99 | 84.37 | 87.51 | 0.7215 |
| LLE | 57 | 81.59 | 82.61 | 80.65 | 0.6349 |
| PCA | 57 | 86.98 | 86.62 | 87.32 | 0.7412 |
| TSVD | 57 | 86.06 | 85.77 | 86.33 | 0.7243 |
| SE | 57 | 84.76 | 85.14 | 84.41 | 0.6970 |
| LASSO | 57 | **88.45** | 88.38 | 88.50 | **0.7711** |

**Fig. 3.**
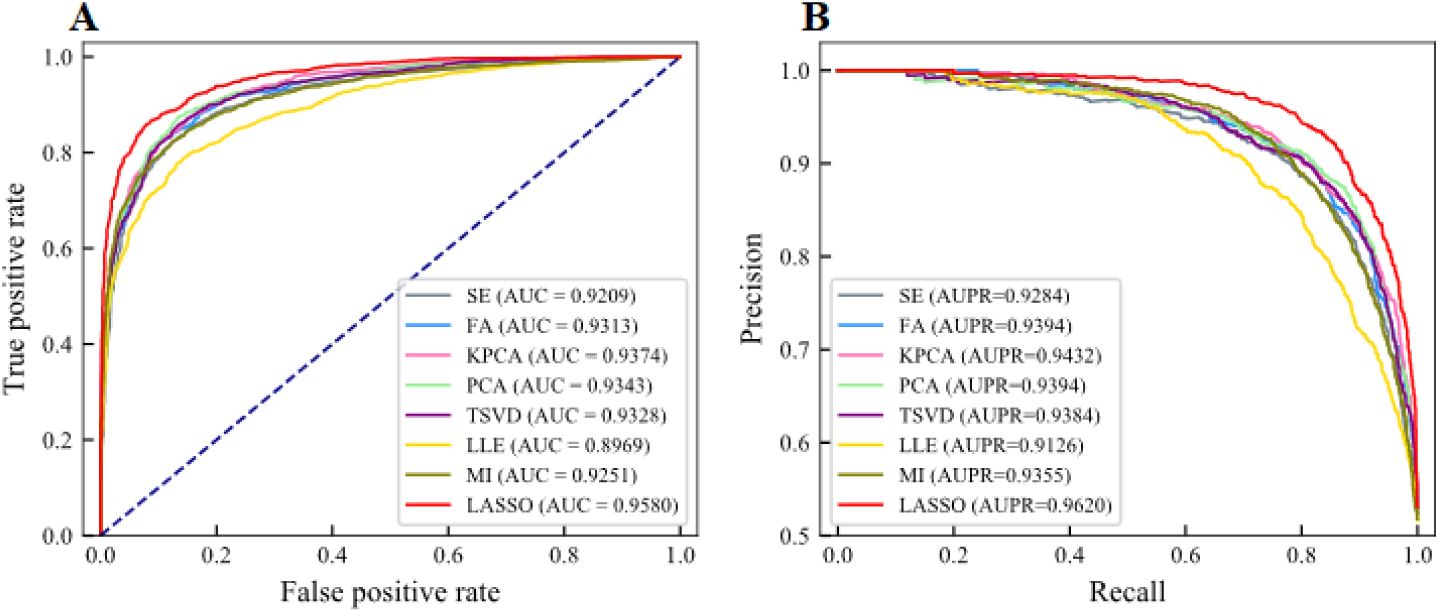
The AUC and AUPR of eight feature extraction methods on the training dataset. (A) ROC curve of eight feature extraction methods. (B) PR curve of eight feature extraction methods.

Table 2 shows that the Acc of MI, FA, KPCA, LLE, PCA, TSVD, SE and LASSO are 84.63%, 85.28%, 85.99%, 81.59%, 86.98%, 86.06%, 84.76% and 88.45%, respectively. The MCC of MI, FA, KPCA, LLE, PCA, TSVD, SE and LASSO are 0.6980, 0.7072, 0.7215, 0.6349, 0.7412, 0.7243, 0.6970 and 0.7711, respectively. The Acc and MCC of LASSO are 1.47%-6.89% and 2.99%-13.62% higher than other feature selection methods, respectively. Therefore, the best feature subset can be obtained by LASSO.

Fig. 3 shows that AUC of MI, FA, KPCA, LLE, PCA, TSVD and SE are 0.9251, 0.9313, 0.9374, 0.8969, 0.9343, 0.9328 and 0.9209, respectively. The AUPR of MI, FA, KPCA, LLE, PCA, TSVD and SE are 0.9355, 0.9394, 0.9432, 0.9126, 0.9394, 0.9384 and 0.9284, respectively. The AUC of LASSO is 0.9580, which is 2.06%-6.11% higher than other methods. The AUPR of LASSO is 0.9620, which is 1.88%-4.94% higher than other methods. It shows that LASSO can eliminate redundant features more effectively than other methods.

### 3.4. Influence of classifier on prediction results

The selection of classification models with strong generalization ability is also the key to build efficient fertility-related proteins prediction models. By comparing prediction results obtained by random forest (RF) [52, 53], k-nearest neighbor (KNN) [54], gradient boosting decision tree (GBDT) [44, 55], Naïve Bayes [56], adaptive boosting (AdaBoost) [57], multi-layer perceptron (MLP) [58], SVM [59] and LightGBM, we choose the best classification algorithm. All classifiers use default parameters and SVM uses polynomial kernel function. The optimal feature subset chose by LASSO on the training dataset is input into different classifiers, respectively. The results are shown in Table 3 and Fig. 4.

**Table 3.** The prediction results of different classifiers on the training dataset.

| Classifier | Acc (%) | Sen (%) | Spe (%) | MCC |
| --- | --- | --- | --- | --- |
| RF | 85.55 | 81.76 | 89.10 | 0.7140 |
| GBDT | 88.00 | 87.89 | 88.11 | 0.7627 |
| Naïve Bayes | 86.74 | 87.75 | 85.80 | 0.7372 |
| KNN | 85.55 | 92.61 | 78.93 | 0.7217 |
| AdaBoost | 87.42 | 88.73 | 86.19 | 0.7520 |
| MLP | 86.16 | 86.13 | 86.19 | 0.7250 |
| SVM | 88.00 | 88.94 | 87.12 | 0.7635 |
| LightGBM | **88.45** | 88.38 | 88.50 | **0.7711** |

**Fig. 4.**
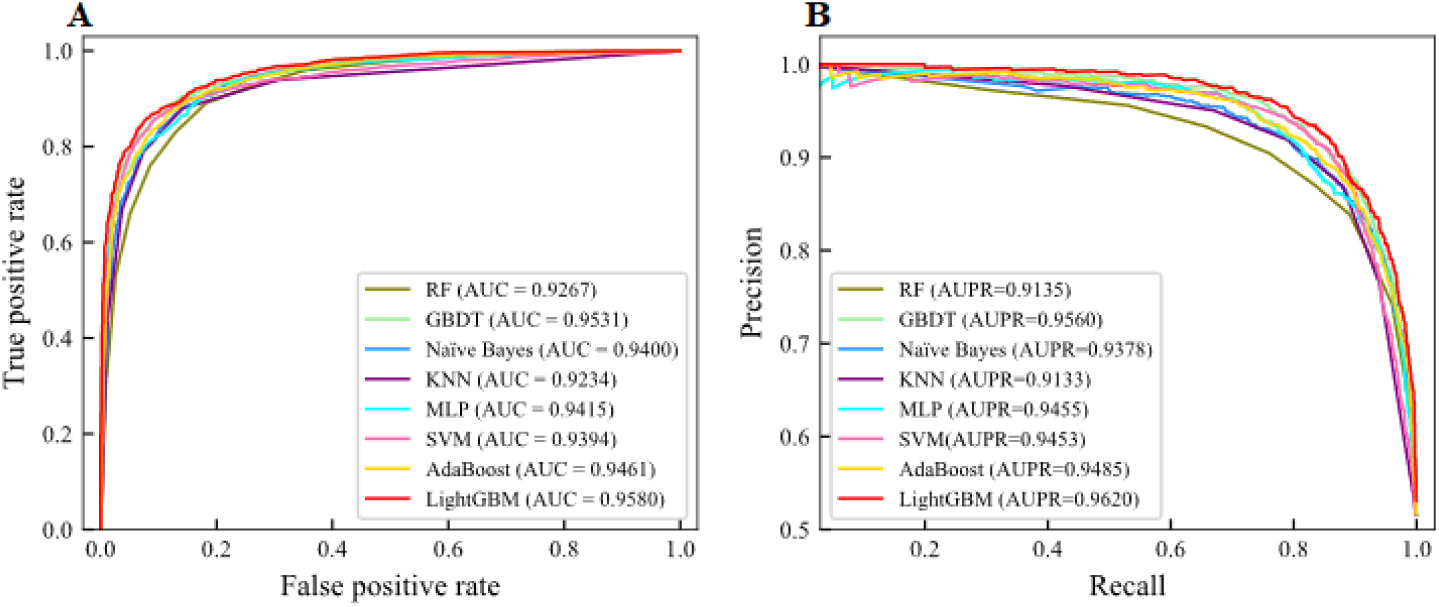
The AUC and AUPR of eight classifiers on the training dataset. (A) the ROC curve of eight classifiers. (B) PR curve of eight classifiers.

Table 3 shows that the Acc of RF, GBDT, Naïve Bayes, KNN, AdaBoost, MLP, SVM and LightGBM are 85.55%, 88.00%, 86.74%, 85.55%, 87.42%, 86.16%, 88.00% and 88.45%, respectively. The MCC of different classifiers are 0.7140, 0.7627, 0.7372, 0.7217, 0.7520, 0.7250, 0.7635 and 0.7711, respectively. The Acc and MCC of LightGBM are the highest, which are 0.45%-2.90% and 0.76%-5.71% higher than those of others.

Fig. 4 shows that on the training dataset, the AUC of LightGBM is 0.9580, which is 3.13%, 0.49%, 1.80%, 3.46%, 1.65%, 1.86% and 1.19% higher than RF, GBDT, Naïve Bayes, KNN, MLP, SVM and AdaBoost. Similarly, the PR curve of LightGBM also enclose PR curves of other classifiers. The AUPR of RF, GBDT, Naïve Bayes, KNN, MLP, SVM and AdaBoost are 0.9135, 0.9560, 0.9378, 0.9133, 0.9455, 0.9453 and 0.9485, respectively. The AUPR of LightGBM is 0.9620, which is 1.00%-4.87% higher than others.

It is proved that LightGBM has better robustness by analyzing the prediction indicators such as Acc, Sen, Spe, MCC, AUC and AUPR on the training dataset of different classifiers. Therefore, we choose LightGBM as the best classifier.

### 3.5. Comparison with existing models

To prove the effectiveness of our model, the Fertility-LightGBM predictions are compared with PrESOgenesis [14] and Fertility-GRU [15]. The specific prediction results are shown in Fig. S2 and Table S6. The prediction results of the training dataset based on the 5-fold cross-validation are shown in Table 4.

**Table 4.** Comparison of prediction results with existing models on training dataset.

| Models | Acc (%) | Sen (%) | Spe (%) | MCC |
| --- | --- | --- | --- | --- |
| PrESOGenesis [14] | 83.0 | 83.6 | 83.0 | 0.67 |
| Fertility-GRU [15] | 85.8 | 88.6 | 83.3 | 0.72 |
| Fertility-LightGBM | **88.5** | 88.4 | 88.5 | **0.77** |

Table 4 shows that the Fertility-LightGBM obtains 88.5% of Acc and 0.77 of MCC through prediction. The Acc of Fertility-LightGBM is 5.5% higher than Acc of PrESOgenesis and 2.7% higher than Acc of Fertility-GRU. The MCC predicted by Fertility-LightGBM is 10.0% higher than that predicted by PrESOgenesis and 5.0% higher than that predicted by Fertility-GRU. Therefore, Fertility-LightGBM has obvious advantages on the training dataset.

We test the generalization ability of the Fertility-LightGBM by the independent dataset test. The Acc, Sen, Spe and MCC are also used as evaluation indicators. The comparison results of PrESOgenesis [14] and Fertility-GRU [15] are shown in Fig. 5.

**Fig. 5.**
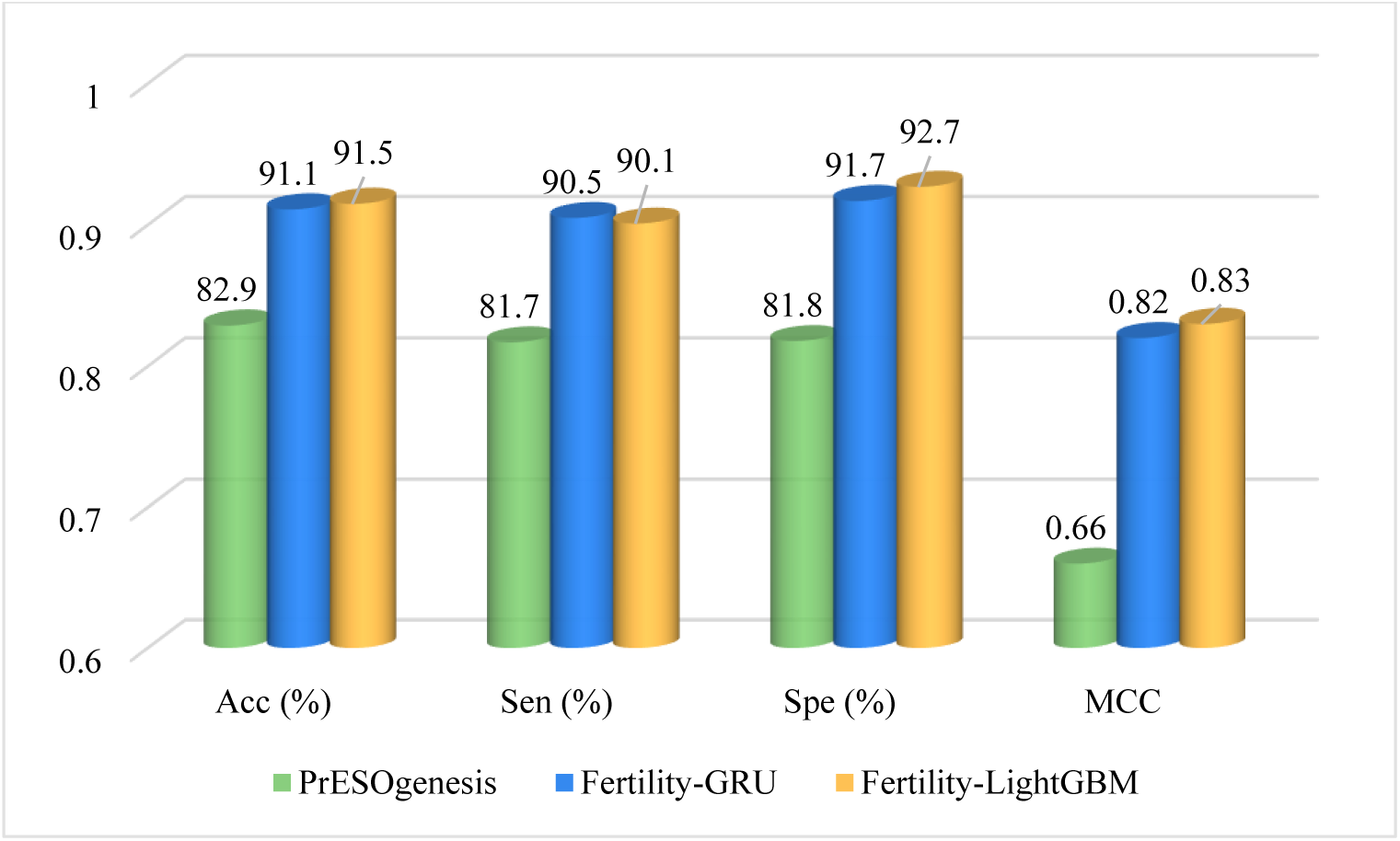
The prediction results on the independent test dataset.

Fig. 5 shows that the Acc of Fertility-LightGBM is 91.5%, which is 0.4%-8.6% higher than those from other models. The MCC of Fertility-LightGBM is 17.0% and 1.0% higher than that of PrESOgenesis and Fertility-GRU, respectively. To sum up, on the independent test dataset, our model improves the accuracy of fertility-related proteins prediction, and the prediction results achieve the desired results.

## 4. Conclusion

Researchers can identify fertility-related proteins to understand their mechanisms, that may deter fertility-related diseases. The construction of prediction models is of great significance for the study of fertility-related proteins. We propose a new prediction model based on LightGBM named Fertility-LightGBM. Multi-information fusion is used to construct the initial feature space which contains physicochemical property information, sequence information and evolutionary information. LASSO is used to delete redundant features in the initial feature space. The LASSO algorithm optimizes the objective function to compress variables with correlation less than the threshold to 0 and eliminate them, so as to achieve the purpose of feature selection. The selected optimal feature subset is used as input information of LightGBM classifier. LightGBM optimizes the sampling method of sample points by GOSS algorithm and compresses the feature dimension by EFB when choosing split points. Compared with traditional machine learning methods, LightGBM supports efficient parallelism and optimize support for category features. Meanwhile, it has the advantage of high efficiency. Fertility-LightGBM can effectively distinguish between fertility-related proteins and non-fertility-related proteins, and it reduces other predictive costs while pushing the research of fertility-related proteins to a new stage of development. Although Fertility-LightGBM can accurately predict fertility-related proteins, there is still much space for improvement. In the future, we will combine proteins structure information and deep learning knowledge to build a more ideal and reliable prediction model of fertility-related proteins.

## Acknowledgments

This work was supported by the National Nature Science Foundation of China (No. 61863010), the Key Research and Development Program of Shandong Province of China (No. 2019GGX101001), and the Natural Science Foundation of Shandong Province of China (No. ZR2018MC007).

## References

[1] G. Anifandis, C. Messini, K. Dafopoulos, S. Sotiriou, I. Messinis, Molecular and cellular mechanisms of sperm-oocyte interactions opinions relative to in vitro fertilization (IVF), Int. J. Mol. Sci. 15 (2014) 12972–12997.

[2] J. Johnson, J. Canning, T. Kaneko, J.K. Pru, J.L. Tilly, Germline stem cells and follicular renewal in the postnatal mammalian ovary, Nature 428 (2004) 145–150.

[3] A. Rodriguez, S.A. Pangas, Regulation of germ cell function by SUMOylation, Cell Tissue Res. 363 (2016) 47–55.

[4] J. Johnson, J. Bagley, M.E. Skaznikwikiel, H. Lee, G.B. Adams, Y. Niikura, K.S. Tschudy, J.C. Tilly, M.L. Cortes, R. Forkert, T. Spitzer, J. Iacomini, D.T. Scadden, J.L. Tilly, Oocyte generation in adult mammalian ovaries by putative germ cells in bone marrow and peripheral blood, Cell 122 (2005) 303–315.

[5] G. Yoshizaki, S. Lee, Production of live fish derived from frozen germ cells via germ cell transplantation, Stem Cell Res. 29 (2018) 103–110.

[6] Y. Park, W. Kwon, S. Oh, M. Pang, Fertility-related proteomic profiling bull Spermatozoa Separated by Percoll, J. Proteome Res. 11 (2012) 4162–4168.

[7] O. D’Amours, G. Frenette, S. Bourassa, É. Calvo, P. Blondin, R. Sullivan, Proteomic markers of functional sperm population in bovines: comparison of low- and high-density spermatozoa following cryopreservation, J. Proteome Res. 17 (2018) 177–188.

[8] J. Schumacher, S. Ramljak, A.R. Asif, M. Schaffrath, H. Zischler, H. Herlyn, Evolutionary conservation of mammalian sperm proteins associates with overall, not tyrosine, phosphorylation in human spermatozoa, J. Proteome Res. 12 (2013) 5370–5382.

[9] A.A. Moura, H. Koc, D.A. Chapman, G.J. Killian, Identification of proteins in the accessory sex gland fluid associated with fertility indexes of dairy bulls: a proteomic approach, J. Androl. 27 (2006) 201–211.

[10] J. Chen, J. Li, Z. You, L. Liu, J. Liang, Y. Ma, M. Chen, H. Zhang, Z. Jiang, B. Zhong, Proteome analysis of silkworm, bombyx mori, larval gonads: characterization of proteins involved in sexual dimorphism and gametogenesis, J. Proteome Res. 12 (2013) 2422–2438.

[11] W. Kwon, S. Rahman, J. Lee, J. Kim, S. Yoon, Y. Park, Y. You, S. Hwang, M. Pang, A comprehensive proteomic approach to identifying capacitation related proteins in boar spermatozoa, BMC Genomics 15 (2014) 897.

[12] C. Légaré, A. Droit, F. Fournier, S. Bourassa, A. Force, F. Cloutier, R. Tremblay, R. Sullivan, Investigation of male infertility using quantitative comparative proteomics, J. Proteome Res. 13 (2014) 5403–5414.

[13] M. Rahimi, M.R. Bakhtiarizadeh, A. Mohammadi-Sangcheshmeh, OOgenesis_Pred: a sequence-based method for predicting oogenesis proteins by six different modes of Chou’s pseudo amino acid composition, J. Theor. Biol. 414 (2017) 128–136.

[14] M.R. Bakhtiarizadeh, M. Rahimi, A. Mohammadi-Sangcheshmeh, V. Shariati J, S.A. Salami, PrESOgenesis: A two-layer multi-label predictor for identifying fertility-related proteins using support vector machine and pseudo amino acid composition approach, Sci. Rep. 8 (2018) 9025.

[15] N.Q.K. Le, Fertility-GRU: identifying fertility-related proteins by incorporating deep-gated recurrent units and original position-specific scoring matrix profiles, J. Proteome Res. 18 (2019) 3503–3511.

[16] L. Fu, B. Niu, Z. Zhu, S. Wu, W. Li, CD-HIT: accelerated for clustering the next-generation sequencing data, Bioinformatics 28 (2012) 3150–3152.

[17] K. Chou, H. Shen, MemType-2L: a web server for predicting membrane proteins and their types by incorporating evolution information through Pse-PSSM, Biochem. Bioph. Res. Co. 360 (2007) 339–345.

[18] W. Qiu, S. Li, X. Cui, Z. Yu, M. Wang, J. Du, Y. Peng, B. Yu, Predicting protein submitochondrial locations by incorporating the pseudo-position specific scoring matrix into the general Chou’s pseudo-amino acid composition, J. Theor. Biol. 450 (2018) 86–103.

[19] H. Shi, S. Liu, J. Chen, X. Li, Q. Ma, B. Yu, Predicting drug-target interactions using Lasso with random forest based on evolutionary information and chemical structure, Genomics 111 (2019) 1839–1852.

[20] B. Yu, S. Li, W. Qiu, M. Wang, J. Du, Y. Zhang, X. Chen, Prediction of subcellular location of apoptosis proteins by incorporating PsePSSM and DCCA coefficient based on LFDA dimensionality reduction, BMC Genomics 19 (2018) 478.

[21] R. Yang, C. Zhang, L. Zhang, R. Gao, A two-step feature selection method to predict cancerlectins by multiview features and synthetic minority oversampling technique, Biomed Res. Int. 2018 (2018) 9364182.

[22] T. Oda, K. Lim, K. Tomii, Simple adjustment of the sequence weight algorithm remarkably enhances PSI-BLAST performance, BMC Bioinformatics 18 (2017) 288.

[23] D.T. Jones, Protein secondary structure prediction based on position-specific scoring matrices, J. Mol. Bio. 292 (1999) 195–202.

[24] B. Manavalan, T.H. Shin, G. Lee, PVP-SVM: sequence-based prediction of phage virion proteins using a support vector machine, Front. Microbiol. 9 (2018) 476.

[25] P. Feng, H. Ding, W. Chen, H. Lin, Naïve Bayes classifier with feature selection to identify phage virion proteins, Comput. Math. Method. M. 2013 (2013) 530696.

[26] H. Nakashima, K. Nishikawa, Discrimination of intracellular and extracellular proteins using amino acid composition and residue-pair frequencies, J. Mol. Bio. 238 (1994) 54–61.

[27] M.S. Khan, M. Hayat, S.A. Khan, N. Iqbal, Unb-DPC: identify mycobacterial membrane protein types by incorporating un-biased dipeptide composition into Chou’s general PseAAC, J. Theor. Biol. 415 (2017) 13–19.

[28] K. Ahmad, M. Waris, M. Hayat, Prediction of protein submitochondrial locations by incorporating dipeptide composition into Chou’s general pseudo amino acid composition, J. Membrane Biol. 249 (2016) 293–304.

[29] H. Zhou, C. Chen, M. Wang, Q. Ma, B. Yu, Predicting Golgi-Resident protein types using conditional covariance minimization with XGBoost based on multiple features fusion, IEEE Access 7 (2019) 144154–144164.

[30] Z. You, L. Zhu, C. Zheng, H. Yu, S. Deng, Z. Ji, Prediction of protein-protein interactions from amino acid sequences using a novel multi-scale continuous and discontinuous feature set, Bioinformatics 15 (2014) S9.

[31] M.N. Davies, A. Secker, A.A. Freitas, E.B. Clark, J. Timmis, D.R. Flower, Optimizing amino acid groupings for GPCR classification, Bioinformatics 24 (2008) 1980–1986.

[32] Z. Chen, P. Zhao, F. Li, A. Leier, T.T. Marquez-Lago, Y. Wang, G.I. Webb, A.I. Smith, R.J. Daly, K.C. Chou, J. Song, iFeature: a Python package and web server for features extraction and selection from protein and peptide sequences, Bioinformatics 34 (2018) 2499–2502.

[33] S. Kawashima, P. Pokarowski, M. Pokarowska, A. Kolinski, T. Katayama, M. Kanehisa, AAindex: amino acid index database, progress report 2008, Nucleic Acids Res. 36 (2007) 202–205.

[34] Z. Zhang, Z. Wang, Z. Zhang, Y. Wang, A novel method for apoptosis protein subcellular localization prediction combining encoding based on grouped weight and support vector machine, Febs Lett. 580 (2006) 6169–6174.

[35] X. Wang, B. Yu, A. Ma, C. Chen, B. Liu, Q. Ma, Protein-protein interaction sites prediction by ensemble random forests with synthetic minority oversampling technique, Bioinformatics 35 (2019) 2395–2402.

[36] B. Tian, X. Wu, C. Chen, W. Qiu, Q. Ma, B. Yu, Predicting protein-protein interactions by fusing various Chou’s pseudo components and using wavelet denoising approach, J. Theor. Biol. 462 (2019) 329–346.

[37] B. Yu, W. Qiu, C. Chen, A. Ma, J. Jiang, H. Zhou, Q. Ma, SubMito-XGBoost: predicting protein submitochondrial localization by fusing multiple feature information and eXtreme gradient boosting, Bioinformatics 36 (2019) 1074–1081.

[38] B. Yu, Z. Yu, C. Chen, A. Ma, B. Liu, B. Tian, Q. Ma, DNNAce: Prediction of prokaryote lysine acetylation sites through deep neural networks with multi-information fusion, Chemometr. Intell. Lab. 200 (2020)103999.

[39] H. Zou, The adaptive lasso and its oracle properties, J. Am. Stat. Assoc. 101 (2006) 1418–1429.

[40] X. Cui, Z. Yu, B. Yu, M. Wang, B. Tian, Q. Ma, UbiSitePred: a novel method for improving the accuracy of ubiquitination sites prediction by using LASSO to select the optimal Chou’s pseudo components, Chemometr. Intell. Lab. 184 (2019) 28–43.

[41] Z. Zhan, Z. You, L. Li, Y. Zhou, H. Yi, Accurate prediction of ncRNA-protein interactions from the integration of sequence and evolutionary information, Front. Genet. 9 (2018) 458.

[42] C. Chen, Q. Zhang, Q. Ma, B. Yu, LightGBM-PPI: Predicting protein-protein interactions through LightGBM with multi-information fusion, Chemometr. Intell. Lab. 191 (2019) 54–64.

[43] G. Ke, Q. Meng, T. Finley, T. Wang, W. Chen, W. Ma, Q. Ye, T. Liu, LightGBM: a highly efficient gradient boosting decision tree, Advances in Neural Information Processing Systems 30 (2017) 3149–3157.

[44] J.H. Friedman, Greedy function approximation: a gradient boosting machine,Ann. Stat. 29 (2001) 1189–1232.

[45] O.P. Tabbaa, C. Jayaprakash, Mutual information and the fidelity of response of gene regulatory models, Phys. Biol. 11 (2014) 046004.

[46] D.A. Engemann, A. Gramfort, Automated model selection in covariance estimation and spatial whitening of MEG and EEG signals, NeuroImage 108 (2015) 328–342.

[47] J. Li, X. Li, D. Tao, KPCA for semantic object extraction in images, Pattern Recogn. 41 (2008) 3244–3250.

[48] X. Liu, D. Tosun, M.W. Weiner, N. Schuff, Locally linear embedding (LLE) for MRI based Alzheimer’s disease classification, NeuroImage 83 (2013) 148–157.

[49] X.Q. Ru, L.D. Wang, L.H. Li, H. Ding, X.C. Ye, Q. Zou, Exploration of the correlation between GPCRs and drugs based on a learning to rank algorithm, Comput. Biol. Med. 119 (2020) 103660.

[50] P. Gao, J. Rong, H. Pu, T. Liu, W. Zhang, X. Zhang, H. Lu, Sparse view cone beam X-ray luminescence tomography based on truncated singular value decomposition, Opt. Express 26 (2018) 23233–23250.

[51] Y. Bengio, O. Delalleau, N.L. Roux, J. Paiement, P. Vincent, M. Ouimet, Learning eigenfunctions links spectral embedding and kernel PCA, Neural Comput. 16 (2004) 2197–2219.

[52] X. Sun, T. Jin, C. Chen, X. Cui, Q. Ma, B. Yu, RBPro-RF: use Chou’s 5-steps rule to predict RNA-binding proteins via random forest with elastic net, Chemometr. Intell. Lab. 197 (2020) 103919.

[53] M. Wang, L. Yue, X. Cui, C. Chen, H. Zhou, Q. Ma, B. Yu, Prediction of extracellular matrix proteins by fusing multiple feature information, elastic net, and random forest algorithm, Mathematics 8 (2020) 169.

[54] K. Chou, H. Shen, Predicting eukaryotic protein subcellular location by fusing optimized evidence-theoretic k-nearest neighbor classifiers, J. Proteome Res. 5 (2006) 1888–1897.

[55] M. Wang, X. Cui, B. Yu, C. Chen, Q. Ma, H. Zhou, SulSite-GTB: identification of protein S-sulfenylation sites by fusing multiple feature information and gradient tree boosting, Neural Comput. Appl. 32 (2020) 13843–13862.

[56] X. Chen, M. Chen, K. Ning, BNArray: an R package for constructing gene regulatory networks from microarray data by using Bayesian network, Bioinformatics 22 (2006) 2952–2954.

[57] I. Mukherjee, C. Rudin, R.E. Schapire, The rate of convergence of AdaBoost, J. Mach. Learn. Res. 14 (2013) 2315–2347.

[58] S.K. Pal, S. Mitra, Multilayer perceptron, fuzzy sets, and classification, IEEE Trans. Neural Netw. 3 (1992) 683–697.

[59] B. Yu, S. Li, C. Chen, J. Xu, W. Qiu, X. Wu, R. Chen, Prediction subcellular localization of Gram-negative bacterial proteins by support vector machine using wavelet denoising and Chou’s pseudo amino acid composition, Chemometr. Intell. Lab. 167 (2017) 102–112.

