## Supplementary Tables, Supplementary Figures for "Fertility-LightGBM: A fertility-related protein prediction model by multi-information fusion and light gradient boosting machine"

#### Table of Contents


#### 1. Supplementary Method Illustration

**Si1.** Encoding based on grouped weigh.

**Si2.** Feature selection.

**Si3.** Machine learning.

**Si4.** Model evaluation.

#### 2. Supplementary Tables

**Table S1.** 20 amino acids are grouped into 7 categories according to the dipole and the volume of the side chain.

**Table S2.** 7 Physicochemical properties of 20 natural amino acids.

**Table S3.** The prediction results of different $\xi$ values on the training dataset.

**Table S4.** The prediction results of different $lag$ values on the training dataset.

**Table S5.** The prediction results of different $L$ values on the training dataset.

**Table S6.** Comparison of prediction results with other models on independent test dataset.

#### 3. Supplementary Figures

**Fig. S1.** Schematic diagram of segmented local descriptor coding construction.

**Fig. S2.** The prediction results on the training dataset.

### 1. Supplementary Method Illustration

##### Si1. Encoding based on grouped weigh .

Different amino acids have different physicochemical properties. Therefore, amino acids can be divided into:

neutral and non-polarity residue $K_1 = \{A, F, G, I, L, M, P, V, W\}$ ,

neutral and polarity residue $K_2 = \{C, N, Q, S, T, Y\}$ ,

acidic residue $K_3 = \{D, E\}$ ,

basic residue $K_4 = \{H, K, R\}$ .

By combining two non-intersect groups, three new partition methods are obtained:

$\{K_{1,} K_2\}$ and $\{K_{3,} K_4\}$ ,

$\{K_{1,} K_3\}$ and $\{K_{2,} K_4\}$ ,

$\{K_{1,} K_4\}$ and $\{K_{2,} K_3\}$ .

##### Si2. Feature selection.

In (11), the regularization coefficient $\delta$ controls the penalty degree of sparse coefficient estimation, $\|w\|_1$ is the $\ell_1$-norm. $w$ , which is the sparse solution, means that in the initial features, only the features of the non-zero component of $w$ will appear in the final model. $\delta \geq 0$ is an adjustable parameter. When the $\delta$ value is large enough, some variable coefficients with low correlation will be compressed to $0$ , then be deleted, so as to achieve the purpose of dimensionality reduction. When the $\delta$ value is small enough, the regularization constraint has little effect on the coefficient, and all attributes will be selected at this time.

##### Si3. Machine learning.

Gradient-based one-side sampling (GOSS) can eliminate a large part of the data with very small gradient, and only use the remaining data to estimate the information gain, so as to avoid the influence of the low gradient long tail part. Since the data with large gradient is more important for information gain, GOSS sets a gradient threshold to retain the samples whose gradient is greater than the threshold, and other samples with small gradient are randomly sampled in proportion. GOSS uses this method to process the data.

##### Si4. Model evaluation.

In the 5-fold cross-validation test, the training dataset is randomly divided into five mutually exclusive subsets with similar size, one of which is the test sample and the other is the model training samples. The cross-validation process is repeated five times and the mean of the 5-fold cross-validation is the result of the classifier performance verification.

In Eq. (11) - Eq. (16), $S_{fertility}$ consists of all positive samples (fertility-related proteins), and $S_{non-fertility}$ consists of all negative samples (non-fertility-related proteins). $\mathrm{TN}$ is the negative samples that are correctly predicted. $\mathrm{TP}$ is the positive samples that are correctly predicted. $\mathrm{FN}$ is the negative samples that are predicted errors. $\mathrm{FP}$ is the positive samples that are predicted errors.

#### 2. Supplementary Tables

**Table S1**

20 amino acids are grouped into 7 categories according to the dipole and the volume of the side chain.

| Group 1 | Group 2 | Group 3 | Group 4 | Group 5 | Group 6 | Group 7 |
|---|---|---|---|---|---|---|
| A, G, V | C | M, S, T, Y | F, I, L, P | H, N, Q, W | K, R | D, E |

**Table S2**

7 Physicochemical properties of 20 natural amino acids.

| Amino acid code | $H_1$ | $H_2$ | VSC | $P_1$ | $P_2$ | SASA | NCISC |
|---|---|---|---|---|---|---|---|
| A | 0.62 | -0.5 | 27.5 | 8.1 | 0.046 | 1.181 | 0.007187 |
| C | 0.29 | -1 | 44.6 | 5.5 | 0.128 | 1.461 | -0.03661 |
| D | -0.9 | 3 | 40 | 13 | 0.105 | 1.587 | -0.02382 |
| E | -0.74 | 3 | 62 | 12.3 | 0.151 | 1.862 | 0.006802 |
| F | 1.19 | -2.5 | 115.5 | 5.2 | 0.29 | 2.228 | 0.03755 |
| G | 0.48 | 0 | 0 | 9 | 0 | 0.881 | 0.1791 |
| H | -0.4 | -0.5 | 79 | 10.4 | 0.23 | 2.025 | -0.01069 |
| I | 1.38 | -1.8 | 93.5 | 5.2 | 0.186 | 1.81 | 0.02163 |
| K | -1.5 | 3 | 100 | 11.3 | 0.219 | 2.258 | 0.01771 |
| L | 1.06 | -1.8 | 93.5 | 4.9 | 0.186 | 1.931 | 0.05167 |
| M | 0.64 | -1.3 | 94.1 | 5.7 | 0.221 | 2.034 | 0.002683 |
| N | -0.78 | 2 | 58.7 | 11.6 | 0.134 | 1.655 | 0.005392 |
| P | 0.12 | 0 | 41.9 | 8 | 0.131 | 1.468 | 0.2395 |
| Q | -0.85 | 0.2 | 80.7 | 10.5 | 0.18 | 1.932 | 0.04921 |
| R | -2.53 | 3 | 105 | 10.5 | 0.291 | 2.56 | 0.04359 |
| S | -0.18 | 0.3 | 29.3 | 9.2 | 0.062 | 1.298 | 0.004627 |
| T | -0.05 | -0.4 | 51.3 | 8.6 | 0.108 | 1.525 | 0.003352 |
| V | 1.08 | -1.5 | 71.5 | 5.9 | 0.14 | 1.645 | 0.057 |
| W | 0.81 | -3.4 | 145.5 | 5.4 | 0.409 | 2.663 | 0.03798 |
| Y | 0.26 | -2.3 | 117.3 | 6.2 | 0.298 | 2.368 | 0.0236 |

*Note:* hydrophobicity ($H_1$), hydrophilicity ($H_2$), volume of side chains, (VSC), polarity ($P_1$), polarizability ($P_2$), solvent accessible surface area (SASA), net charge index of side chains, (NCISC).

**Table S3**

The prediction results of different $\xi$ values on the training dataset.

| Parameter | Acc (%) | Sen (%) | Spe (%) | MCC(%) |
|---|---|---|---|---|
| $\xi$=1 | 83.54 | 86.34 | 80.91 | 67.56 |
| $\xi$=2 | 84.08 | 86.90 | 81.44 | 68.64 |
| $\xi$=3 | 84.39 | 87.46 | 81.50 | 69.23 |
| $\xi$=4 | 83.91 | 87.25 | 80.78 | 68.36 |
| $\xi$=5 | 83.81 | 86.41 | 81.37 | 68.20 |

| | | | | |
|---|---|---|---|---|
| $\xi$=6 | 84.08 | 87.46 | 80.91 | 68.81 |
| $\xi$=7 | 84.73 | 87.54 | 82.10 | 69.99 |
| $\xi$=8 | 84.08 | 87.25 | 81.11 | 68.74 |
| $\xi$=9 | **84.83** | 87.04 | 82.76 | **70.09** |
| $\xi$=10 | 84.49 | 87.89 | 81.30 | 69.50 |
| $\xi$=11 | 84.32 | 87.32 | 81.50 | 69.20 |
| $\xi$=12 | 84.46 | 87.68 | 81.44 | 69.50 |
| $\xi$=13 | 84.32 | 87.82 | 81.04 | 69.30 |
| $\xi$=14 | 84.53 | 88.24 | 81.04 | 69.70 |
| $\xi$=15 | 83.71 | 87.32 | 80.31 | 68.80 |
| $\xi$=16 | 84.12 | 87.61 | 80.84 | 68.80 |
| $\xi$=17 | 84.46 | 88.03 | 81.11 | 69.50 |
| $\xi$=18 | 84.05 | 87.46 | 80.84 | 68.60 |
| $\xi$=19 | 83.61 | 86.90 | 80.51 | 67.80 |
| $\xi$=20 | 84.18 | 87.82 | 80.78 | 69.00 |
| $\xi$=21 | 84.22 | 87.89 | 80.78 | 69.02 |
| $\xi$=22 | 84.42 | 87.96 | 81.11 | 69.43 |
| $\xi$=23 | 84.32 | 87.54 | 81.30 | 69.18 |
| $\xi$=24 | 84.25 | 87.25 | 81.44 | 69.03 |
| $\xi$=25 | 83.81 | 86.97 | 80.84 | 68.20 |
| $\xi$=26 | 83.88 | 87.11 | 80.84 | 68.29 |
| $\xi$=27 | 83.44 | 86.48 | 80.58 | 67.40 |
| $\xi$=28 | 83.78 | 87.11 | 80.64 | 68.09 |
| $\xi$=29 | 83.50 | 87.04 | 80.18 | 67.57 |
| $\xi$=30 | 83.26 | 86.62 | 80.11 | 67.10 |
| $\xi$=31 | 83.71 | 86.97 | 80.64 | 67.99 |
| $\xi$=32 | 83.78 | 86.90 | 80.84 | 68.03 |
| $\xi$=33 | 83.91 | 87.25 | 80.78 | 68.32 |
| $\xi$=34 | 84.18 | 87.46 | 81.11 | 68.94 |
| $\xi$=35 | 83.74 | 86.90 | 80.78 | 68.00 |
| $\xi$=36 | 83.50 | 86.69 | 80.51 | 67.53 |
| $\xi$=37 | 84.32 | 87.82 | 81.04 | 69.20 |
| $\xi$=38 | 84.05 | 87.25 | 81.04 | 68.60 |
| $\xi$=39 | 84.25 | 87.61 | 81.11 | 69.05 |
| $\xi$=40 | 84.08 | 87.96 | 80.45 | 68.80 |
| $\xi$=41 | 84.01 | 87.32 | 80.91 | 68.59 |
| $\xi$=42 | 83.74 | 86.90 | 80.78 | 68.02 |
| $\xi$=43 | 84.22 | 87.46 | 81.17 | 68.96 |
| $\xi$=44 | 84.25 | 87.89 | 80.84 | 69.04 |
| $\xi$=45 | 84.01 | 86.83 | 81.37 | 68.50 |
| $\xi$=46 | 83.98 | 87.54 | 80.64 | 68.58 |
| $\xi$=47 | 84.12 | 87.46 | 80.97 | 68.76 |
| $\xi$=48 | 83.54 | 86.55 | 80.71 | 67.60 |
| $\xi$=49 | 83.98 | 87.32 | 80.84 | 68.52 |
| $\xi$=50 | 83.91 | 87.39 | 80.64 | 68.40 |

**Table S4**

The prediction results of different *lag* values on the training dataset.

| Parameter | Acc (%) | Sen (%) | Spe (%) | MCC(%) |
|---|---|---|---|---|
| *lag* =1 | 70.14 | 69.30 | 70.93 | 40.33 |
| *lag* =2 | 72.32 | 69.93 | 74.57 | 44.57 |
| *lag* =3 | 72.49 | 69.51 | 75.30 | 44.97 |
| *lag* =4 | 73.76 | 71.41 | 75.95 | 47.48 |
| *lag* =5 | 74.81 | 71.97 | 77.47 | 49.60 |
| *lag* =6 | 74.98 | 71.76 | 78.00 | 49.93 |
| *lag* =7 | 75.22 | 72.61 | 77.67 | 50.37 |
| *lag* =8 | 74.30 | 71.41 | 77.01 | 48.55 |
| *lag* =9 | 75.08 | 72.39 | 77.61 | 50.14 |
| *lag* =10 | 74.98 | 72.04 | 77.74 | 49.90 |
| *lag* =11 | 75.36 | 72.18 | 78.33 | 50.67 |
| *lag* =12 | 75.53 | 71.83 | 78.99 | 51.02 |
| *lag* =13 | 75.19 | 71.90 | 78.27 | 50.33 |
| *lag* =14 | 75.43 | 72.11 | 78.53 | 50.80 |
| *lag* =15 | 75.63 | 71.76 | 79.26 | 51.30 |
| *lag* =16 | 75.77 | 72.89 | 78.47 | 51.52 |
| *lag* =17 | 76.11 | 73.45 | 78.60 | 52.20 |
| *lag* =18 | 76.18 | 72.39 | 79.72 | 52.34 |
| *lag* =19 | 75.32 | 72.18 | 78.27 | 50.64 |
| *lag* =20 | 75.94 | 70.77 | 80.78 | 51.90 |
| *lag* =21 | 75.87 | 71.83 | 79.65 | 51.80 |
| *lag* =22 | 75.97 | 72.25 | 79.45 | 52.00 |
| *lag* =23 | **76.55** | 72.04 | 80.78 | **53.20** |
| *lag* =24 | 76.21 | 71.97 | 80.18 | 52.30 |
| *lag* =25 | 75.80 | 70.92 | 80.38 | 51.70 |
| *lag* =26 | 76.35 | 72.96 | 79.52 | 52.70 |
| *lag* =27 | 76.45 | 72.96 | 79.72 | 53.00 |
| *lag* =28 | 75.90 | 71.62 | 79.92 | 51.80 |
| *lag* =29 | 75.39 | 71.27 | 79.26 | 50.80 |
| *lag* =30 | 75.94 | 72.25 | 79.39 | 51.90 |
| *lag* =31 | 75.90 | 71.48 | 80.05 | 51.84 |
| *lag* =32 | 75.73 | 71.90 | 79.32 | 51.45 |
| *lag* =33 | 75.56 | 71.55 | 79.32 | 51.15 |
| *lag* =34 | 75.39 | 71.55 | 78.99 | 50.76 |
| *lag* =35 | 75.63 | 70.77 | 80.18 | 51.30 |
| *lag* =36 | 75.63 | 71.48 | 79.52 | 51.28 |
| *lag* =37 | 75.32 | 71.06 | 79.33 | 50.69 |
| *lag* =38 | 75.36 | 71.62 | 78.86 | 50.75 |
| *lag* =39 | 75.26 | 71.13 | 79.13 | 50.57 |
| *lag* =40 | 75.19 | 70.63 | 79.46 | 50.40 |
| *lag* =41 | 75.53 | 71.76 | 79.06 | 51.05 |
| *lag* =42 | 75.80 | 72.11 | 79.26 | 51.60 |
| *lag* =43 | 75.66 | 71.06 | 79.98 | 51.34 |
| *lag* =44 | 75.63 | 70.70 | 80.25 | 51.31 |
| *lag* =45 | 75.77 | 71.83 | 79.46 | 51.50 |

| | | | | |
|---|---|---|---|---|
| *lag* =46 | 75.80 | 71.48 | 79.85 | 51.60 |
| *lag* =47 | 76.14 | 71.55 | 80.45 | 52.28 |
| *lag* =48 | 75.60 | 70.77 | 80.12 | 51.23 |
| *lag* =49 | 75.36 | 70.28 | 80.12 | 50.75 |
| *lag* =50 | 75.53 | 71.06 | 79.72 | 51.10 |

**Table S5**

The prediction results of different $L$ values on the training dataset.

| Parameter | Acc (%) | Sen (%) | Spe (%) | MCC(%) |
|---|---|---|---|---|
| $L$ =1 | 66.63 | 68.38 | 64.99 | 33.44 |
| $L$ =2 | 68.51 | 71.06 | 66.11 | 37.26 |
| $L$ =3 | 67.59 | 69.79 | 65.52 | 35.46 |
| $L$ =4 | 68.23 | 68.87 | 67.64 | 36.60 |
| $L$ =5 | 67.86 | 68.52 | 67.24 | 35.90 |
| $L$ =6 | 68.41 | 68.45 | 68.36 | 36.95 |
| $L$ =7 | 68.03 | 70.00 | 66.18 | 36.23 |
| $L$ =8 | 68.85 | 71.06 | 66.78 | 37.90 |
| $L$ =9 | 67.31 | 68.80 | 65.92 | 34.77 |
| $L$ =10 | 68.78 | 69.65 | 67.97 | 37.70 |
| $L$ =11 | 68.98 | 71.20 | 66.91 | 38.21 |
| $L$ =12 | 69.46 | 69.93 | 69.02 | 39.00 |
| $L$ =13 | 69.56 | 70.77 | 68.43 | 39.26 |
| $L$ =14 | 69.22 | 70.21 | 68.30 | 38.57 |
| $L$ =15 | 68.64 | 70.14 | 67.24 | 37.50 |
| $L$ =16 | 70.35 | 71.76 | 69.02 | 40.90 |
| $L$ =17 | 68.51 | 69.01 | 68.03 | 37.13 |
| $L$ =18 | 69.63 | 69.86 | 69.42 | 39.35 |
| $L$ =19 | 68.58 | 69.15 | 68.03 | 37.24 |
| $L$ =20 | 68.23 | 69.44 | 67.11 | 36.70 |
| $L$ =21 | 69.63 | 70.77 | 68.56 | 39.40 |
| $L$ =22 | 68.00 | 69.44 | 66.64 | 36.17 |
| $L$ =23 | 68.78 | 69.65 | 67.96 | 37.68 |
| $L$ =24 | 68.58 | 69.15 | 68.03 | 37.30 |
| $L$ =25 | 68.88 | 69.15 | 68.63 | 37.80 |
| $L$ =26 | 70.79 | 71.90 | 69.75 | 41.72 |
| $L$ =27 | 69.73 | 70.77 | 68.76 | 39.64 |
| $L$ =28 | 69.43 | 70.99 | 67.97 | 39.03 |
| $L$ =29 | 70.31 | 71.48 | 69.22 | 40.78 |
| $L$ =30 | 69.33 | 70.28 | 68.43 | 38.80 |
| $L$ =31 | 69.02 | 69.23 | 68.82 | 38.16 |
| $L$ =32 | 69.05 | 69.86 | 68.29 | 38.26 |
| $L$ =33 | 69.70 | 70.21 | 69.22 | 39.47 |
| $L$ =34 | 69.33 | 69.15 | 69.48 | 38.72 |
| $L$ =35 | 69.84 | 70.63 | 69.09 | 39.80 |
| $L$ =36 | 69.97 | 70.00 | 69.94 | 40.05 |
| $L$ =37 | 69.36 | 69.86 | 68.89 | 38.83 |
| $L$ =38 | 70.18 | 71.83 | 68.63 | 40.51 |
| $L$ =39 | 70.08 | 70.77 | 69.42 | 40.26 |

| | | | | |
|---|---|---|---|---|
| $L$ =40 | 69.26 | 70.35 | 68.23 | 38.60 |
| $L$ =41 | 70.01 | 71.62 | 68.49 | 40.19 |
| $L$ =42 | 69.90 | 70.35 | 69.48 | 39.90 |
| $L$ =43 | 70.28 | 71.62 | 69.02 | 40.69 |
| $L$ =44 | 71.13 | 71.69 | 70.61 | 42.37 |
| $L$ =45 | 70.31 | 71.27 | 69.42 | 40.80 |
| $L$ =46 | 70.35 | 71.20 | 69.55 | 40.90 |
| $L$ =47 | 69.84 | 70.49 | 69.22 | 39.90 |
| $L$ =48 | 70.55 | 70.99 | 70.14 | 41.30 |
| $L$ =49 | **71.23** | 72.25 | 70.28 | **42.60** |
| $L$ =50 | 70.31 | 70.56 | 70.08 | 40.70 |

**Table S6**

Comparison of prediction results with other models on independent test dataset.

| Models | Acc (%) | Sen (%) | Spe (%) | MCC |
|---|---|---|---|---|
| PrESOgenesis | 82.9 | 81.7 | 81.8 | 0.66 |
| Fertility-GRU | 91.1 | 90.5 | 91.7 | 0.82 |
| Fertility-LightGBM | **91.5** | 90.1 | 92.7 | **0.83** |

#### 3. Supplementary Figure

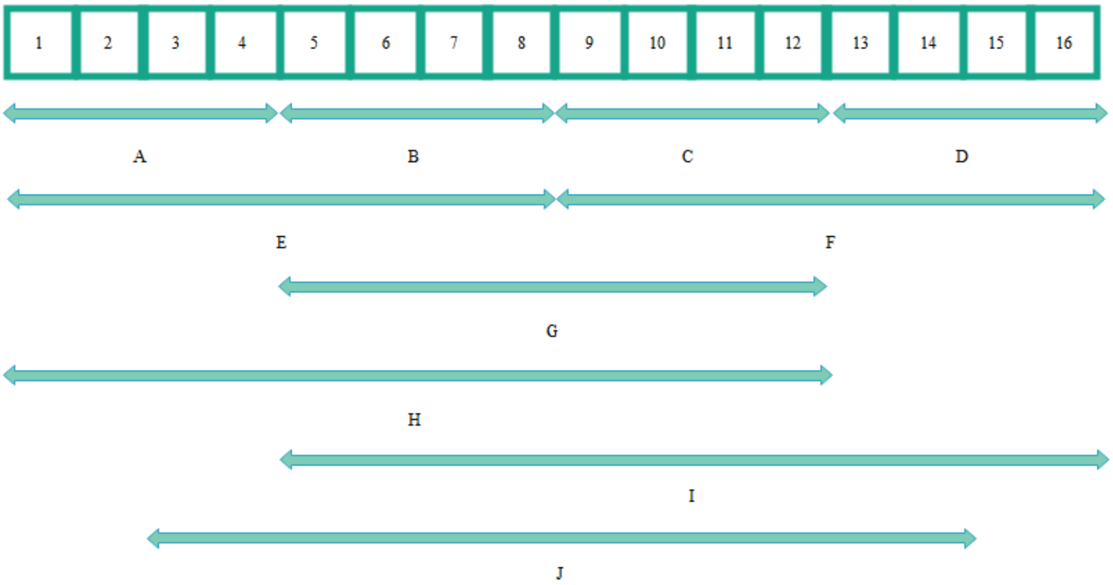


**Fig. S1.** Schematic diagram of segmented local descriptor coding construction.

*Note:* The areas (A-D) and (E-F) are generated by dividing the entire protein sequence into quarters and halves, respectively. Regions G, H, I and J represent the 50%, 75%, 75% and 75% fragments of the entire protein sequence, respectively.

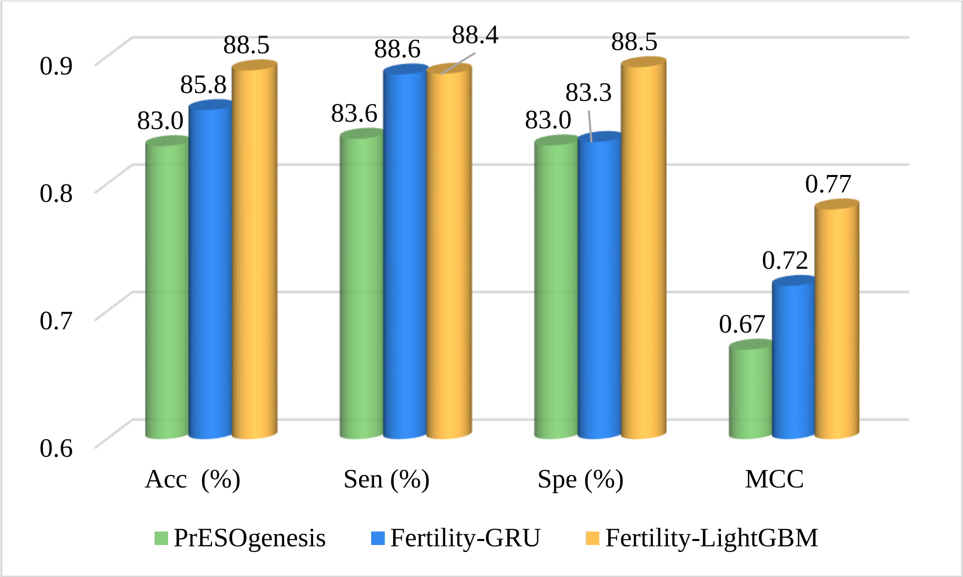


**Fig. S2.** The prediction results on the training dataset.
